# Metagenomic sequencing for rapid identification of *Xylella fastidiosa* from leaf samples

**DOI:** 10.1101/2021.05.12.443947

**Authors:** Veronica Roman-Reyna, Enora Dupas, Sophie Cesbron, Guido Marchi, Sara Campigli, Mary Ann Hansen, Elizabeth Bush, Melanie Prarat, Katherine Shiplett, Melanie L. Lewis Ivey, Joy Pierzynski, Sally A. Miller, Francesca Peduto Hand, Marie-Agnes Jacques, Jonathan M. Jacobs

**Affiliations:** Department of Plant Pathology, The Ohio State University, Columbus, OH, USA; Infectious Disease Institute, The Ohio State University, Columbus, OH, USA; Univ Angers, Institut Agro, INRAE, IRHS, SFR QUASAV, F-49000 Angers, France; French Agency for Food, Environmental and Occupational Health & Safety, Plant Health Laboratory, Angers, France; Department of Agriculture, Food, Environment and Forestry, University of Florence, Italy; School of Plant and Environmental Sciences, Virginia Tech, Blacksburg, VA, USA; Animal Disease Diagnostic Laboratory, Ohio Department of Agriculture, Reynoldsburg, OH, USA; Department of Plant Pathology, The Ohio State University, Wooster, OH, USA; C. Wayne Ellett Plant and Pest Diagnostic Clinic, Department of Plant Pathology, The Ohio State University Reynoldsburg, OH, USA

**Keywords:** *Xylella fastidiosa*, metagenomics, diagnostics, short-read sequencing

## Abstract

*Xylella fastidiosa* (*Xf*) is a globally distributed plant pathogenic bacterium. The primary control strategy for *Xf* diseases is eradicating infected plants; therefore, timely and accurate detection is necessary to prevent crop losses and further pathogen dispersal. Conventional *Xf* diagnostics primarily relies on quantitative PCR (qPCR) assays. However, these methods do not consider new or emerging variants due to pathogen genetic recombination and sensitivity limitations. We developed and tested a metagenomics pipeline using in-house short-read sequencing as a complementary approach for affordable, fast, and highly accurate *Xf* detection. We used metagenomics to identify *Xf* to strain level in single and mixed infected plant samples at concentrations as low as one picogram of bacterial DNA per gram of tissue. We also tested naturally infected samples from various plant species originating from Europe and the United States. We identified *Xf* subspecies in samples previously considered inconclusive with real-time PCR (Cq > 35). Overall, we showed the versatility of the pipeline by using different plant hosts and DNA extraction methods. Our pipeline provides taxonomic and functional information for *Xf* diagnostics without extensive knowledge of the disease. We hope this pipeline can be used for early detection of *Xf* and incorporated as a tool to inform disease management strategies.

**IMPORTANCE:** *Xylella fastidiosa (Xf)* destructive outbreaks in Europe highlight this pathogen’s capacity to expand its host range and geographical distribution. The current disease diagnostic approaches are limited by a multiple-step process, biases to known sequences, and detection limits. We developed a low-cost, user-friendly metagenomic sequencing tool for *Xf* detection. In less than three days, we were able to identify *Xf* subspecies and strains in field-collected samples. Overall, our pipeline is a diagnostics tool that could be easily extended to other plant-pathogen interactions and implemented for emerging plant threat surveillance.

## INTRODUCTION

*Xylella fastidiosa* (*Xf*), is a globally distributed insect-transmitted plant pathogenic bacterium, causing diseases on a large hosts range. To date, 595 plant species grouped belonging to 85 botanical families have been reported as *Xf* hosts (1), some of which are of major socio-economic interest, such as grapevine, olive, citrus, coffee and almond (2). *Xf* colonizes the xylem vessels of plants where it forms biofilms (3) that, together with tyloses and gums produced by the plant in response to the infection (4), limit water translocation. Infected hosts display symptoms of leaf scorches and plant dieback finally followed by plant death (3).

*Xf* was first described in and limited to the Americas but recently emerged in Europe, highlighting the pathogen’s capacity to expand its host range and geographical distribution (2, 5). The pathogen was reported in Italy in 2013, where is currently devastating Apulian olive production, then detected in France in 2015, Spain in 2016 and Portugal in 2018, both on cultivated as well as spontaneous Mediterranean plant species (2). The primary control strategy for *Xf* diseases includes eradication of hosts; therefore, fast and accurate detection is necessary to prevent major losses to growers and further pathogen dispersal.

The diagnostic of diseases caused by fastidious pathogens like *Xf* is difficult. This difficulty is increased as infected plants may remain asymptomatic for very long periods of time, which is associated with low bacterial concentrations, and by an irregular distribution of the pathogens in the plants (6). It is of major interest to develop reliable and highly sensitive tools for detection and detailed identification that can be used directly on plant extracts. Current standards for *Xf* diagnostics primarily rely on quantitative real-time PCR (qPCR) assays to detect and sometimes identify the bacterium (7–12), followed by the amplification and sequencing of two, for subspecies identification, to seven housekeeping genes (*cysG, gltT, holC, leuA, malF, nuoL* and *petC*) for is Sequence Type (ST) determination and phylogeny reconstruction (2) (Fig. 1A). Five subspecies are proposed in *X. fastidiosa*, ie. *fastidiosa*, *multiplex*, *pauca*, *morus*, and *sandyi* (13–15). However, whole genome analyses revealed similarities of the subspecies *fastidiosa*, *morus* and *sandyi*, which cluster into one clade. Moreover, genome analysis indicated high frequency of horizontal gene transfer and recombination among *Xf* subspecies (14–16).

**Fig. 1.**
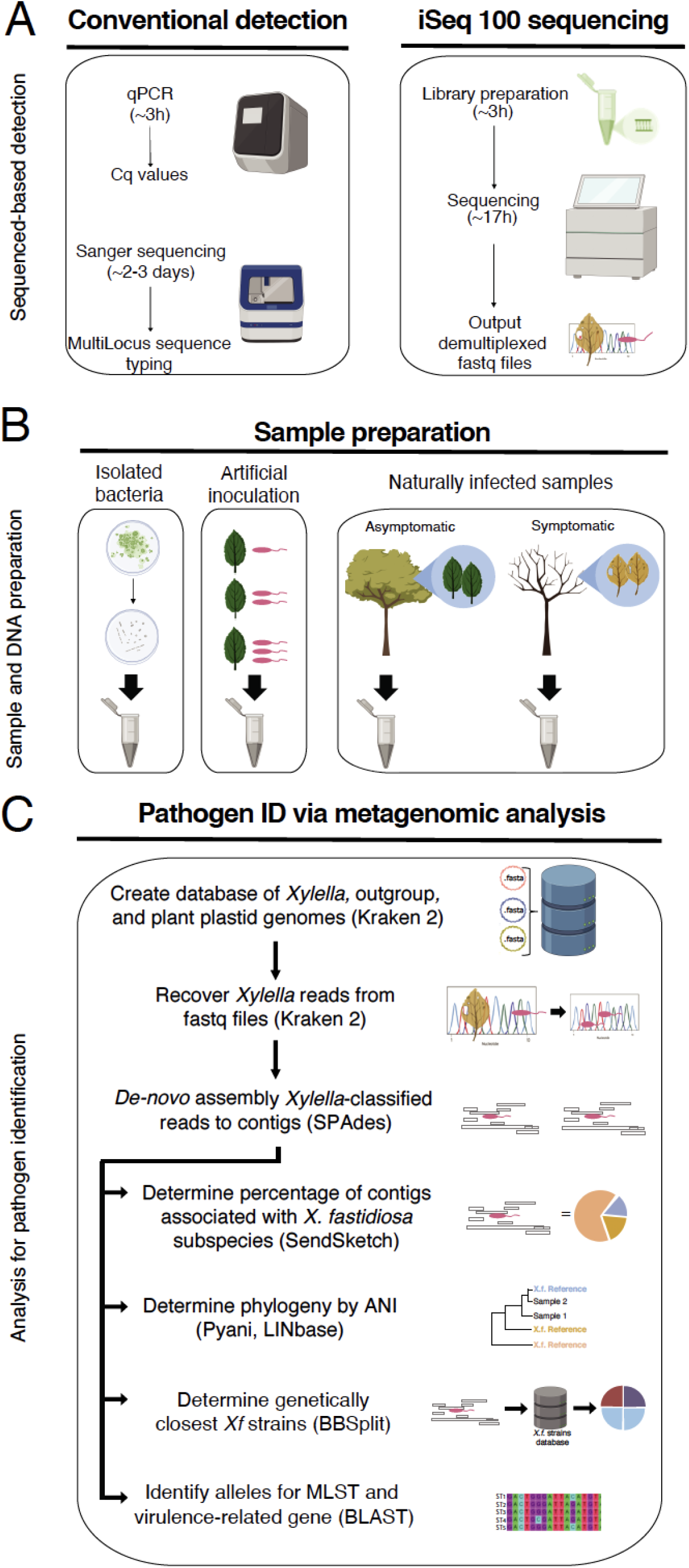
Metagenomics for diagnostic pipeline. **A**) Sequenced-based detection. Two approaches were used for *Xf* detection: conventional detection and iSeq 100 sequencing. For conventional, samples were analyzed using qPCR assays, Harper’s test or tetraplex Dupas’s test, and MLST involving Sanger sequencing of seven housekeeping genes. iSeq 100 libraries were prepared according to manufacturer. After 17h of sequencing, demultiplexed samples were recover from the machine and used for subsequent analysis. B) Sample preparation. The samples used for the pipeline were DNAs extracted from bacterial strains in culture, spiked plant material, and naturally infected samples. **C**) Pathogen identification via metagenomic analysis. Demultiplexed fastq reads from all samples were then used for metagenomic analysis. We created a database to recover *Xf* reads using Kraken2. The database contained *Xylella*, *Xanthomonas* and *Escherichia coli* genomes. We also added plant plastid genomes to remove false positive results. *Xf* reads were recovered from the fastq files. The *Xf* recovered reads were de-novo assembled to obtain *Xf* contigs, using SPADes. The *Xf* contigs were used in four different analyses, subspecies identification, phylogeny reconstruction, identification of the genetically closest strain with a sequenced genome and alleles from specific genes. To determine subspecies, we used the tool SendSketch. To reconstruct phylogeny, we calculated ANI using Pyani and the website tool LINbase (https://linbase.org/). To determine the genetically closest known *Xf* strain, we detected the number of hits to each *Xf* strain using BBSplit. To identify specific genes alleles, we calculated the percentage of identity to the seven MLST genes (*cysG*, *gltT, holC, leuA, malF, nuoL, petC*), and the percentage of similarity to *17* virulence-related proteins using local BLAST+. Graphics were created with BioRender.

Plant samples infected by more than one *Xf* strain belonging to several subspecies are not uncommon and are not easy to detect, (17, 18). Nevertheless, current methods do not consider new or emerging variants resulting from pathogen genetic recombination (14). For example, qPCR with high Cq values (>35) are considered inconclusive (2), making decisions about disease control difficult. A complementary tool for diagnostics is the use of Next-Generation Sequencing (19) (Fig. 1). Because this approach can be directly used on plant extracts, it is not biased towards known sequences and provides more information about the pathogen genome, such as virulence traits. Metagenomics, the study of genetic material from environmental samples, beyond whole genome sequencing allows for the detection of strains from several subspecies and ST at the same time from the host (20). Recently, the use of long-read sequencing as diagnostic tool identified *Xf* subspecies and ST from infected samples (12, 21).

In this study, we developed and tested a metagenomics pipeline using in-house short-read sequencing as a complementary approach for affordable and accurate *Xf* detection. We were able to use metagenomics to identify *Xf* to strain level in single and mixed infected plant samples, at concentrations as low as one picogram of bacterial DNA per gram of tissue. In addition, we tested naturally infected field samples from Europe and the United States. We identified *Xf* subspecies in samples with Cq values equal to and greater than 37, which is beyond the threshold of detection for the standard and certified qPCR methods (2). Overall, we developed a robust diagnostics pipeline that could be easily extended to other pathogens and implemented for surveillance of emerging agricultural threats.

## RESULTS

### Metagenomics for diagnostics pipeline

We developed and tested a metagenomics pipeline for *Xf* detection and subspecies identification (Fig. 1). We tested this pipeline based on three types of DNA samples: from bacterial colonies in culture, spiked plant samples, and naturally infected plant samples (Fig. 1B). To recover and identify *Xf* subspecies and compare it to the already sequenced genomes, we developed a pipeline that uses six different tools and custom-made databases (22) (Fig. 1C). The pipeline recovers *Xf* reads with the software Kraken2 and a custom-made database (22). The database has user-specified genomes for *Xf* reads identification. The user-specified genomes belonged to *Xylella* (n=81), *Xanthomonas* sp. (n=10), *E. coli* (n=1) and several plant sequences from NCBI (Table S1). The database had plant sequences because some *Xf* genomes from NCBI contained plant genomic DNA sequences. We could not clean all 18S sequences, plant plastids, or chloroplast reads from the NCBI *Xf* genomes. Therefore, the plant reads in the database serve as a filter to ensure plant reads were not misidentifying as *Xf* reads.

After Kraken2, the pipeline *de-novo* assembled the recovered *Xf* reads into contigs with the program SPAdes (23). The pipeline used the *Xf* contigs for four different analyses: 1) subspecies identification, 2) phylogeny reconstruction, 3) identification of the already sequenced genetically closest strains, and 4) alleles for MLST profile and virulence-related genes determination. The pipeline used *Xf* contigs, and the tool SendSketch from the BBMap software to identify subspecies. Then it used Pyani and LINbase software to reconstruct phylogeny by ANI (24, 25). Next, it assigned *Xf* strains to each *Xf* contig to identify the closest strains with the tool BBSplit from BBMap software. Finally, to identify specific genes or alleles, the pipeline used local BLAST with two type of subjects: a) one subject was the reported MLST allele genes and b) the other subject was the protein sequences from genes associated with virulence.

### *Xf* identification from *in-silico* prepared samples

To test the pipeline sensitivity, we used *in-silico* samples with target (e.g. *Xf*) and non-target DNAs (e.g. non-host plant). The samples included variable amounts of non-host barley (*Hordeum vulgare*) sequenced reads *in silico* spiked with *Xff* CFBP 7970 reads. We obtained a strong linear correlation between the Kraken2 results and the proportion of spiked *Xf* sequence reads (y =103.21x - 0. 0127; R^2^ = 1) (Fig. S1). We recovered *Xf* reads and assembled them as contigs using SPAdes. With the *Xf* contigs, we performed BLAST analysis to identify MLST alleles and virulence genes. We were able to identify one to four MLST-related gene for samples spiked with 0.5 to 2.4% *Xf* reads (Table S3). This result indicated that we cannot capture the full MLST gene set for ST identification with less than 2.4% *Xf* reads (Table S3). We calculated, for all samples, the percentage of gene similarity to the virulence-related genes (Table S4). The percentage of gene similarity increased with the higher number of spiked *Xf* reads. Samples with a lower number of *Xf* reads had a low genome coverage to recover and analyze complete gene sequences (Table S4, S7).

We then identified *Xf* subspecies using the *Xf* contigs. Since the *in-silico* samples only identified *Xff* reads, we expected that SendSketch assigned all contigs to *Xff*. However, we found that 9 to 15% of *Xff* contigs were instead assigned to *Xfm*. Based on these results, we did two additional analyses to determine the best approach to analyze the *Xf* subspecies composition. For the first analysis, we hypothesized that complete assembled genomes would reduce the percentage of reads assigned to other subspecies. To test this, we created a smaller Kraken2 database with 30 genomes instead of 81. These 30 *Xf* genomes had complete assemblies. We recovered 1 to 2% fewer *Xf* reads with the new database, and the subspecies distribution remained the same (data not shown). The results indicated that *Xf* subspecies classification is not related to the level of genome assembly. Therefore, the original Kraken2 database with 81 NCBI *Xf* genomes was retained for all further analyses.

For the second analysis, we manually assessed the SendSketch sensitivity to mixed infections with new *in-silico* samples. The new samples included a set amount of barley reads *in silico* spiked with variable amounts of *Xff* CFBP 7970 and *Xfm* CFBP8418 reads (Table S2). For these new *in-silico* samples, we recovered *Xf* reads with Kraken2, assembled the reads as contigs and run SendSketch to identify subspecies. When using BLAST, a certain number of *Xf* contigs mapped equally (100% identity) to *Xff* and *Xfm* (*Xf* core contigs). We observed that *Xf* core contigs are directly proportional to the total of *Xf* recovered reads and samples with a higher *Xff* to *Xfm* spiked reads ratio (Table S2, Fig. S2). Moreover, the tool SendSketch randomly assigned the subspecies to *Xf* contigs with 100% identity to *Xff* and *Xfm*. Consequently, we developed a manual correction to separate single from mixed infections. We only used samples with either *Xfm* or *Xff; Xfp* was not part of the analysis. The correction consists of calculating the logarithm of the *Xfm*: *Xff* contigs ratio. Corrected log-ratios from 0.081 to 0.4 are considered a mixed infection. Log-ratios below 0.08 will be a single *Xff* infection, and higher than 0.43 will be *Xfm* single infection (Fig. S2).

To evaluate the pipeline with samples free of *Xf*, we used extracted DNAs of two healthy plant samples and a non-*Xylella* controls (i.e., barley leaves infiltrated with *Xanthomonas*). For the artificially inoculate barley samples, Kraken2 software recovered 20 to 30% of the total reads as *Xanthomonas*. For all four *Xf*-free samples, Kraken2 recovered six to 19 *Xf* reads (Table S2). All these *Xf* reads corresponded to plant reads based on the BLAST webtool from NCBI. Based on these results, the pipeline considers a sample *Xf*-free when it cannot recover more than 19 *Xf* reads (Fig. S3).

### *Xf* identification from isolated bacteria

To test the capacity of iSeq 100 sequencing, we used six gDNAs from isolated bacteria and two known *Xf* gDNAs as control. The six *Xf* gDNAs were isolated from Italian field samples (See Material and Methods). The two control *Xf* gDNA samples were *Xff* CFBP 7970 (CFBP 7970 iSeq100) and *Xfm* CFBP 8418 (CFBP 8418 iSeq100). All eight gDNA samples were sequenced with the iSeq 100 System. For all eight samples, Kraken2 recovered 99% of the total reads as *Xf* reads (Table S2). We assembled the *Xf* reads as contigs and classified them into subspecies. For CFBP 7970 and CFBP 8418 for which a genome was already available, 54-78% of the contigs corresponded to *Xf* core contigs, 44-65% to their respective subspecies, and 2% to the closest subspecies. On average, within the six Italian samples, 20% of the contigs corresponded to *Xf* core contigs, 25% to *Xfm,* and 2% *to Xff*.

The ANI values were consistent with the *Xf* contig abundance (Fig. 2, Table S2). All six Italian samples and CFBP 8418 iSeq100 had 99 to 100% identity to *Xfm* and less than 97% identity to *Xff* and *Xfp*. The control sample, CFBP 7970 iseq100, had 100% identity to *Xff* and less than 98% identity to *Xfm* and *Xfp* (Fig. 2, Table S2).

**Fig. 2:**
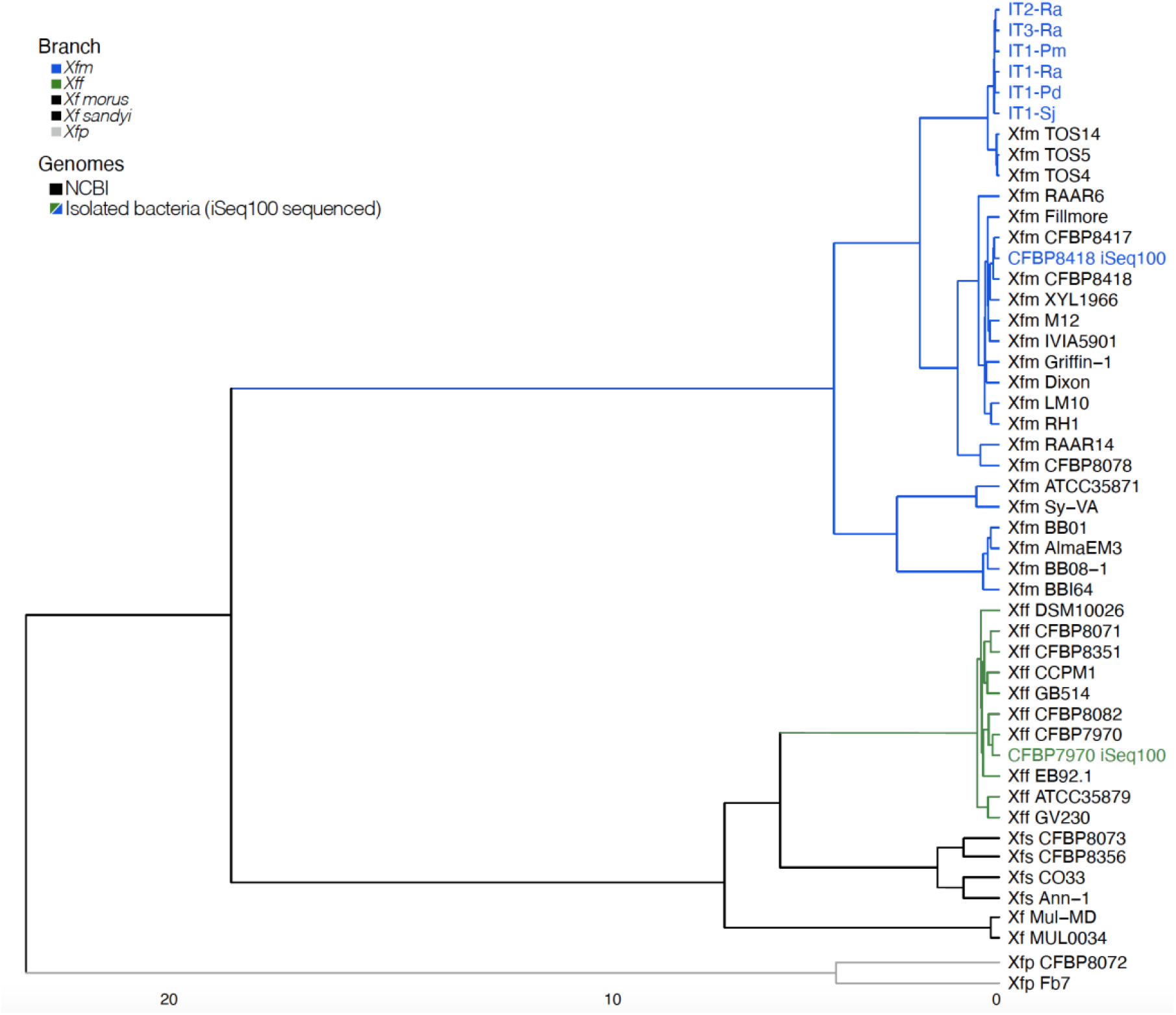
Phylogenetic reconstruction of isolated bacteria used in this study. The cluster analysis is based on average nucleotide identity values from Pyani. Branch colors indicate different *Xf* subspecies: *Xff (green*), *Xfm (blue*), *Xfp (gray*), *Xf* subspecies *morus* and *sandyi* (black). The sequenced gDNA from isolated bacteria are indicated in blue or green. The *Xf* genomes obtained from NCBI are indicated in black. The cluster was plotted using ComplexHeatmap R package.

For strain identification, we used the program BBSplit and the Harvest suite. For each sample, we selected the top three closest strains based on the program BBSplit output. Then, we used these closest strains and the sample to compare the number of single nucleotide polymorphisms (SNPs) with the Harvest suite. For the sample CFBP 8418 iSeq100, the closest strain with 30 SNPs was *Xfm* CFBP 8418 (Table S5, S6). For the sample CFBP 7970 iSeq100, the closest strain with fewer SNPs was *Xff* CFBP 7970. The six Italian samples had the same three closest *Xfm* strains, TOS5, TOS4, and TOS14. All six samples had fewer SNPs when compared against the strain *Xfm* TOS4. The three TOS strains and the Italian samples were isolated from the outbreak area of Monte Argentario, Tuscany, Italy (26).

We performed BLAST analysis to identify MLST alleles and virulence genes for all eight isolated bacteria with the assembled *Xf* contigs. For virulence genes, the sample CFBP 7970 iSeq100 had 100% similarity to all *Xff* virulence genes except for rpfE (96.4%) (Table S4). The sample CFBP 8418 iSeq100 had 100% similarity to all *Xfm* virulence genes except for pilB (99.8%). The six Italian samples had the same similarity percentages for all *Xfm* virulence genes except for hemagglutinin (95.7 to 100%). We were able to identify all virulence genes and to complete the allelic profiles for ST identification (Table S3). As expected, the ST identified for the sample CFBP 7970 iSeq100 was ST2, and the one for CFBP 8418 iSeq100 was ST6. The six Italian samples had the same ST87 number.

### *Xf* identification from spiked plant samples

We tested the pipeline with DNA extracted from grapevine petioles and midribs artificially inoculated with known bacterial concentrations of the strain *Xff* CFBP 7970, *Xfm* CFBP 8418, or an equal mix of both strains. Kraken2 output recovered 0.01% to 74.8% of the total sequences as *Xf* reads (Table S2). The percentage of recovered *Xf* reads had a positive correlation with Log10 CFU values (R^2^= 0.9876) (Fig. 3A) and Cq values (R^2^= 0.9141) (data not shown). The pipeline detected *Xf* with lowest bacterial concentration tested in this study (1× 10^4^ CFU/ml), equivalent to a Cq 28.85 and 1.62 pg.μL^−1^.

**Fig. 3:**
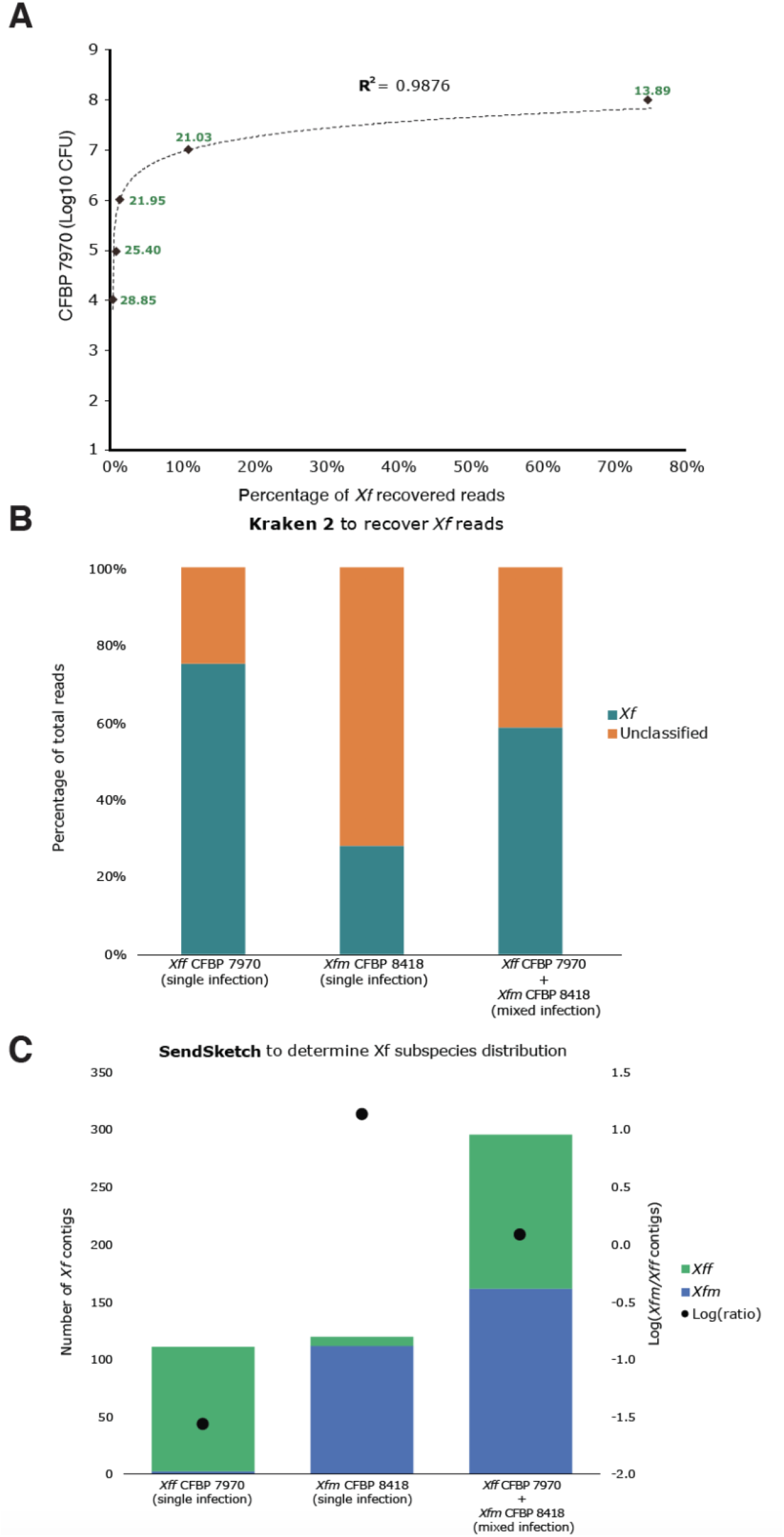
Spiked samples with different dilutions and mixed samples. **A)** scatter plot comparing the Log10 CFU with the percentage of *Xff* CFBP7970 recovered reads from total read number. The dotted line indicates a logarithmic trendline, y = 0. 4222ln(x) + 7. 9511; R^2^=9876. The green numbers indicated the Cq values for each sample. **B)** Percentage of *Xf*-recovered reads by Kraken2 from single (*Xf*f, 1e8; *Xf*m, 1e7 CFU)- and mixed-infected (1e7 CFU) samples. Teal bars indicate *Xf* reads, and orange bars indicate unclassified reads. Unclassified show reads with no similarity to *Xf* reads, like plant or other microorganisms reads. **C)** Proportion of *Xf* subspecies from total *Xf* contigs in single and mixed infections. The black dots indicated the log-ratio as a manual correction to detect single and mixed infection. *Xfm* and *Xff* are indicated in blue and green respectively.

After *Xf* contig assembly, we were able to identify the subspecies for all samples, and the log-ratio separated single from mixed infections (Fig. 3B, 3C). The log-ratio for *Xff*-single infections varied between −1.56 and −0.23. The log-ratio for *Xfm*-single infection was 1.15, while for mixed-strains infection was 0.08. The ANI values confirmed the *Xff* and *Xfm* subspecies for single infected samples (Fig. S4, Table S2). The mixed sample (*Xff* CFBP 7970 + *Xfm* CFBP 8418) had a higher ANI value for *Xfm*. This result was consistent with a higher number of *Xfm* contigs for the mixed-strain sample (Fig. 3C, Table S2).

Based on BBSplit results, the genetically closest strains sequenced for most of the *Xff*-single infections, was CFBP 7970, followed by ATCC 35879, GV230, and TPD4 (Table S5). For *Xfm*-single infection, the genetically closest *Xfm* strains were Dixon and CFBP 8418. For the mixed infection, 90% of *Xf* contigs were assigned to *Xf*m Dixon and 6% to *Xff* strains.

With the assembled *Xf* contigs, we performed BLAST analysis to identify MLST alleles and virulence genes. We identified the ST number for three of the seven artificially inoculated samples. The mixed infected sample of grapevine inoculated with the strain CFBP 8418 were identified as ST6 (Tables S4). The grapevine sample inoculated with CFBP 7970 (10^8^ CFU/ml) was ST2. We could not assign an ST the sample for grapevine sample inoculated with the strain CFBP 7970 (10^7^ CFU/ml) because we only identified six of the seven MLST alleles. In contrast to MLST analysis, we detected at least two virulence genes per sample (Table S4). The sample with the lowest CFU values, CFBP 7970 (10^4^ CFU/ml), had 42% similarity to *Xff* hemagglutinin and 41% similarity to *Xfm* pilQ. For the remaining *Xff*-single infected samples, the percentage of similarity to a single subspecies increased with the higher CFU number, which is also associated with higher genome coverage (Table S7). For the *Xfm*-single infection, all the virulence genes had 100% similarity to *Xfm*. For the mixed infection, all the virulence genes had 100% similarity to *Xfm* and, on average, 98% to *Xff*.

### *Xf* identification from field-collected samples

Finally, we tested the iSeq 100 sequencing capacity with European and American field samples (Table S2). We used 24 samples with Cq values ranging from 21 to 40 based on Harper’s qPCR assay. We used three samples that were negative based on the same qPCR assay. The DNAs from the 27 samples were extracted from six different hosts: *Olea europaea*, *Polygala myrtifolia* (France and Italy), *Quercus ilex*, *Spartium junceum*, *Rhamnus alaternus*, and *Vitis vinifera*.

Kraken2 recovered 0.004% to 1.43% of the total reads as *Xf* (Table S2). We assembled the *Xf* reads into one to 2896 contigs. We found all samples had at least one contig with at least 400 bp. The limit of *Xf* detection with Harper and tetraplex qPCR correspond to 30-37 Cq values (17). Therefore, we evaluated 16 samples that either had less than 30 *Xf* contigs, were classified as *Xf*-negative, or had Cq higher than 30 (2). We used each *Xf* contig from the 16 samples as query for a Nucleotide BLAST search using webtool from NCBI. Eleven of the 16 samples gave 100% identity to *Xf* genomes. Hence, these 11 samples were considered *Xf*-positive. All contigs from the other five samples FR1-Pm, FR1-Oe, IT6-Sj, IT11-Sj, and US1-Vv, had 100% identity to chloroplast and 18S plant sequences but none to *Xf*. Therefore, these five samples were considered *Xf*-negative (Fig. 4). With our pipeline, we were able to detect *Xf* in samples considered inconclusive by qPCR according to Harper’s (EPPO 2019).

**Fig. 4.**
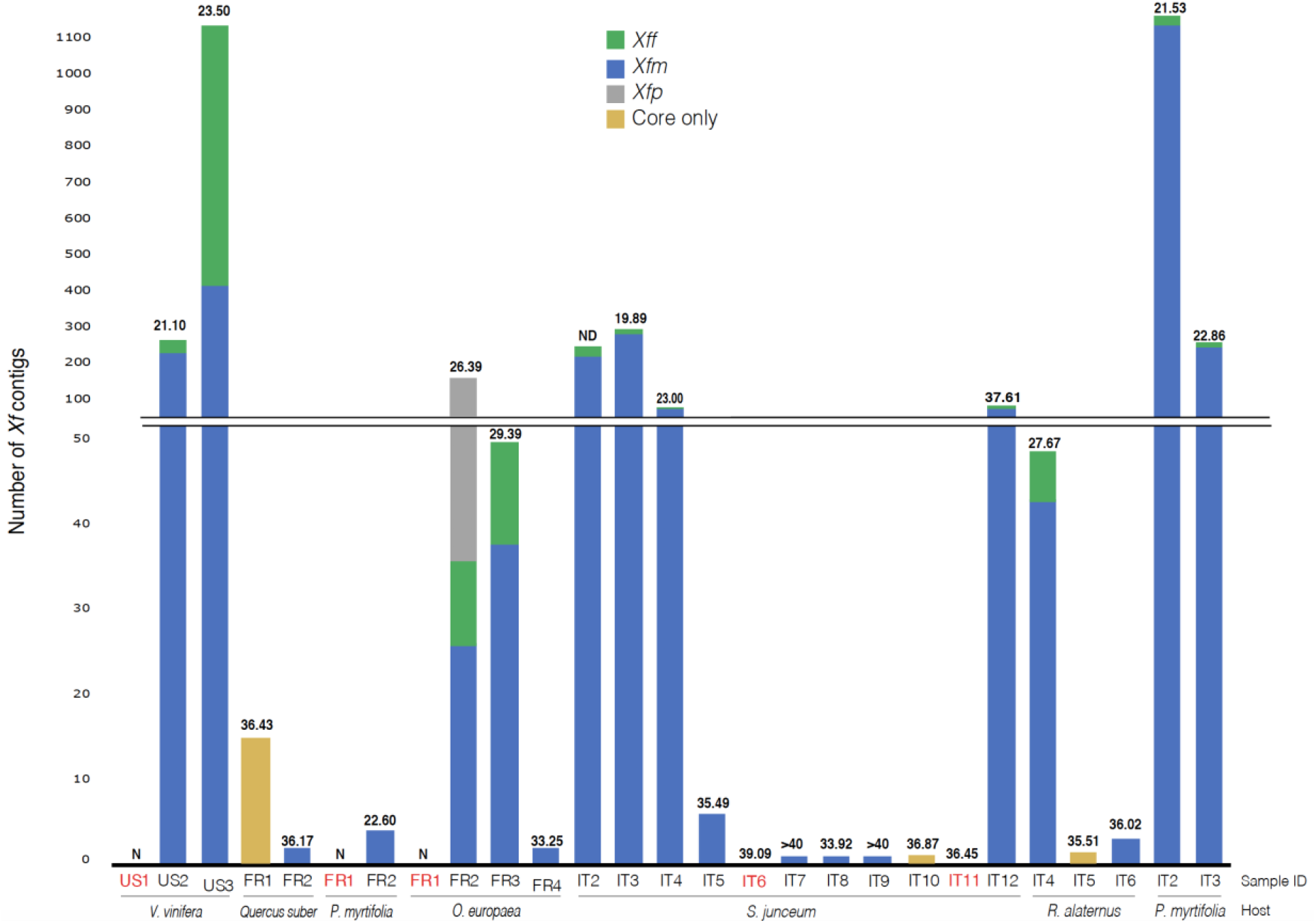
*Xf* from field-collected samples and read mapping to subspecies from database. Stacked columns indicate the number of *Xf* contigs with 100% identify to each *Xf* subspecies. Samples indicated in red were *Xf*-negative with our pipeline. *Xff*: represents the sum *of Xf* subsp. *fastidiosa, Xf morus, and Xf*. *sandyi*; *Xfm*: *Xf* subsp. *multiplex*; *Xfp*: *Xf* subsp. *pauca*. Only Core indicates samples that only have *Xf* core contigs. The Cq values are indicated on the top of each bar. ND is not determined. N is negative for *Xf* based on qPCR Sample code and hosts are indicated on the X-axis. Each country of origin is indicated in the sample ID: France (FR), Italy (IT) and USA (US) along with the host from which they were isolated.

We then used the contigs from the 22 *Xf*-positive samples for subspecies classification. Overall, the samples had 50 to 100% of contigs classified as *Xf* core contigs (Fig. 4). Three French samples (FR2-Qi, FR2-Pm, FR4-Oe) and five Italian samples (IT5-Sj, IT7-Sj, IT8-Sj, IT9-Sj, IT6-Ra) had 1 to 6 contigs assigned as *Xfm*. The French sample FR3-Oe, seven Italian samples (IT2-Sj, IT3-Sj, IT4-Sj, IT12-Sj, IT4-Ra, IT2-Pm, IT3-Pm) and the USA sample US2-Vv, were *Xfm*-single infected based on the manual log correction (log-ratio > 0.43) (Table S2). The sample FR2-Oe had 17% contigs assigned to *Xfp* and the sample US3-Vv was *Xff*-single infected (log-ratio < 0.08).

Then, we determined the ANI values and *Xf* strain composition for the 22 *Xf*-positive samples. Both results were consistent with the subspecies identification. The Italian samples had 99-100% ANI to *Xfm*. Five French samples (FR1-Qi, FR2-Pm, FR3-Oe, FR4-Oe, FR2-Qi) had 99-100% ANI to *Xfm* and FR2-Oe 98% ANI to *Xf*p (Fig. 5A, Table S2). The sample US2-Vv had 99% ANI to *Xfm*, and US3-Vv had 100% to *Xff* (Fig. 5A). For strain distribution, all the French, USA, and three Italian samples (IT2-Sj, IT4-Sj and IT5-Sj) had more than 40% *Xf* contigs had 100% identity to one strain (Table S5, Fig. 5B). Except for IT2-Sj, Italian samples had most of the contigs assigned to the three strains TOS4, TOS5, and TOS14. The sample IT2-Sj had more reads assigned to *Xfm* RAAR14. Overall, the tools SendSketch, Pyani and Bbsplit validated the qPCR subspecies results for field samples.

**Fig. 5:**
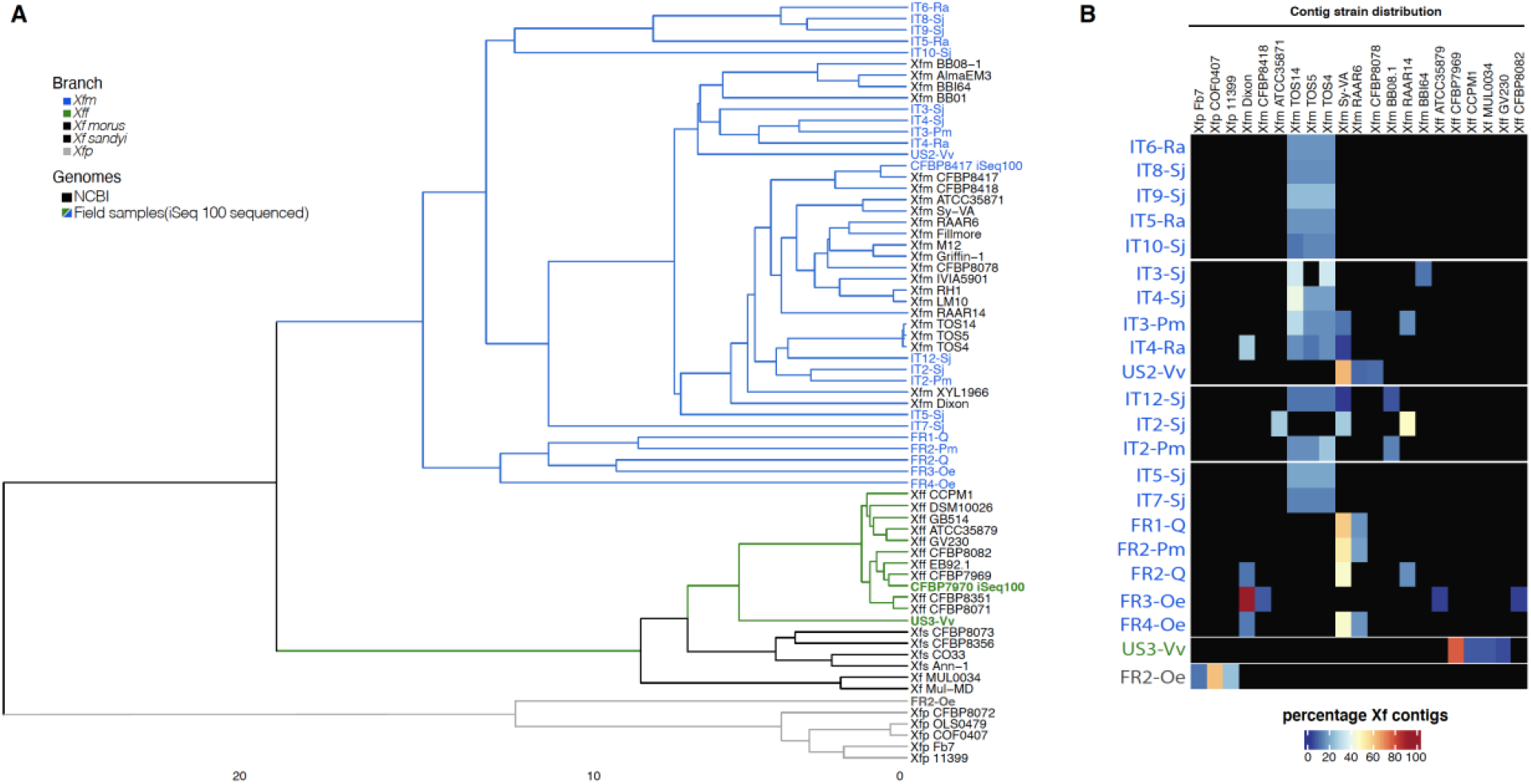
Metagenomics analyses of field samples identifies bacterial subspecies. A) The dendrogram indicates the distance and cluster analysis based on ANI values using NCBI whole genomes and assembled *Xf* contigs. Branch colors represent each *Xf* subspecies. Blue, gray and green names indicate iSeq 100 sequenced samples. B) The heatmap shows the percentage of unique contigs assigned to each *Xf* strain. The cluster and strain distribution were plotted using ComplexHeatmap R package.

We performed BLAST analysis to identify MLST alleles and virulence genes for the 22 infected samples. For 16 of 22 *Xf*-positive samples, we found the percentage of gene similarity to be 26 to 100% for at least one virulence gene (Table S4). Distinct from single gene analysis, we only identified some MLST-related alleles for four samples (US3-Vv, FR2-Oe, IT2-Pm, IT3-Pm); consequently, we could not identify the ST number (Table S3).

To compare some of our results with a high-performance, deep-sequencing Illumina platform as a control, we selected nine samples for re-sequencing with the MiSeq platform using the same iSeq 100 libraries from this study. The nine samples were IT3-Pm, IT5-Sj, FR2-Oe, FR3-Oe, US1-Vv, US2-Vv, FR1-Pm, FR2-Pm, IT9-Sj (Table S8, Fig. S5). We analyzed the MiSeq sequences with our pipeline and recovered 0.005%-0.792% of total reads as *Xf* reads with Kraken2. We assembled the *Xf* reads into contigs and manually assessed all samples with less than 30 *Xf* contigs. The NCBI Blastn analysis indicated that the samples US1-Vv and FR1-Pm, had contigs with 100% identity to plant reads; therefore, we confirmed they were *Xf*-negative samples. The other seven samples were considered *Xf*-positive. We followed the pipeline to identify and determine subspecies, phylogeny, genetically closest sequenced genome strains, MLST profile, and virulence-related genes. The results for subspecies and phylogeny identification were the same between MiSeq and iSeq100 sequencers (Fig. S5), but there were some differences for the other three analysis results (Table S2, Table S8). For the genetically closest sequenced genome strains analysis, all samples gave the same strain distribution as iSeq100 results, except FR2-Oe which showed *Xfp* OLS0478 instead of *Xfp* COF0407 as the most abundant strain. These two *Xfp* strains are phylogenetically close. For MLST analysis, we identified four more alleles for the sample IT3-Pm with the MiSeq platform than with iSeq100 (Table S3), while we only detected two alleles in the sample US2-Vv sequenced with Miseq. We were only able to detect MLST alleles for the sample FR2-Oe with the iSeq100 platform. We were able to calculate the percentage of gene similarity for more virulence genes with the MiSeq platform than with the iSeq100 platform. The variation between Illumina platforms was not consistent.

## DISCUSSION

In this study, we developed a user- friendly metagenomic pipeline to identify and determine *Xylella fastidiosa* subspecies from field-collected samples without the need for pathogen isolation. We demonstrated the flexibility of the pipeline by using seven different plant hosts and three DNA extraction methods. We recovered and assembled *Xf* reads into contigs from total DNA samples. We used percentage of similarity to a single subspecies to identify *Xf* subspecies and validated the results through phylogeny and strain proximity. Finally, we examined potential virulence-related genes among all sequenced samples.

To recover *Xf* reads from field samples with the tool Kraken2, we used *Xf* genomes available on NCBI. We found plant plastid reads in all genomes obtained from pure *Xf* cultures, except for *Xff* CFBP7970. We decided to add plant reads in the database to filter out potential plant contamination. We still recovered *Xf* reads for some plant samples reported as *Xf*-negative. Therefore, we manually examined the contigs of samples with less than 30 *Xf* contigs to determine if they have a low *Xf* concentration or are negative samples. More than the Cq values, the contig evaluation will be necessary to determine if a sample is truly *Xf* -negative.

We observed that the Kraken2 tool not only recovered *Xf* reads but also identified *Xf* subspecies. We decided not to use Kraken2 to identify subspecies because we found that the recovery is affected by incomplete NCBI *Xf* genomes subspecies information. Kraken2 uses by default the NCBI taxonomy to classify reads, if the genomes used to build the database does not have a subspecies information it will keep most of the reads at the species level. To improve the subspecies resolution, we decided to use contigs instead of reads and the tool SendSketch (BBMap tool). Contigs or assembled reads increases the coverage and reduces false-positive reads (27). We used Sendskecth because it uses MinHash algorithm to be fast and it takes into account whole genomes and do not uses taxonomy.

After we identified samples as *Xf*-positive, we defined *Xf* subspecies. We observed that some samples had mapped contigs to both *Xff* and *Xfm*. These are contigs most likely associated with core sequences as only 3% of *Xff* and *Xfm* genomes are different. However, it is also possible to have a percentage of annotation error due to sequencing contamination (28, 29). To determine if the samples were single- or mixed-infected, we corrected the results by calculating the Log *Xfm: Xff* contigs ratio (see Materials and methods). Once we defined the Log values for *Xff, Xfm,* and *Xfm+Xff* (mixed) infections, we validated the presence of single *Xfm* infections in all the tested European samples. Identifying *Xf* subspecies in ornamental and crop plants is essential for the correct application of eradication measures or for plant movements within the EU territory according to regulation (EU) 2020/1201 (30).

The number of reads generated by the iSeq100 sequencer highlighted some limitations of the pipeline. For example, the ST were not determined for any field sample because we could not recover the complete sequences of the seven genes. This is probably associated with low genome coverage. The low number of reads also hampers deep SNP diversity and intersubspecific homologous recombination analyses. Other sequencing systems, with higher reads output than the iseq100, can also be used with this pipeline as the input is fastq files. With a high number of reads, the pipeline will provide better resolution to recover the MSLT genes.

Some of the diagnostic tools for *Xf* diagnostics are qPCR and Sanger sequencing. These tools require amplification of known *Xf* genome regions, but as is the case of MLST, do not consider new or emerging variants. Moreover, these tools introduce bias due to primer design, have unresolved results with high Cq values, and may take longer since they follow a multistep process. Our pipeline complements these conventional tools by obtaining metagenomic data directly from symptomatic or asymptomatic samples and increasing the detection power. We found some discrepancies between the number of recovered *Xf* reads and Cq values. These differences could be caused by PCR inhibitors or the genomic target region for qPCR that underestimates the bacterial concentration (31, 32). Metagenomics sequencing is becoming a more affordable and faster approach for diagnostics. For example, the whole detection/identification with qPCR and MLST scheme could have an estimated cost of 52-54USD per sample and takes three to four days to detect one to seven genes. With the iSeq100, it could cost 50-70USD (when having 12 samples in the same run) and take two days but while also allowing to get a complete genomic analysis of the plant and its pathogenic and commensal microbiota.

In conclusion, our pipeline provides *Xf* taxonomy and functional information for diagnostics without extensive knowledge of the host or pathogen. The pipeline databases used for the analysis can be public repositories or privately collected gDNA and could be adapted by the user and tailored to different plant pathogens. The sequencing can be adapted to be an in-house system as the library preparation and sequencing are user-friendly and not limited by the DNA quality or quantity. The analysis can be adjusted to detect several pathogens simultaneously. We hope this pipeline can be used for early detection of *Xf* or other crop pathogens and incorporated as part of management strategies.

## MATERIAL AND METHODS

### *Xylella* fastidiosa strains

The strain *Xf* subsp. *fastidiosa* (*Xff*) CFBP 7970 isolated in the United States (Florida) in 2013 from *Vitis vinifera* and *Xf* subsp. *multiplex* (*Xfm*) CFBP 8418 isolated in France in 2015 from *Spartium junceum* were provided by the French Collection of Plant-Associated Bacteria (CIRM-CFBP (CIRM-CFBP https://www6.inra.fr/cirm_eng/CFBP-Plant-Associated-Bacteria) and used as controls for whole-genome sequencing. Both strains were cultivated on modified PWG medium (Gelrite 12 g. L^−1^; soytone 4 g. L^−1^; bacto tryptone 1 g. L^−1^; MgSO_4_. 7H_2_O 0, 4 g. L^−1^; K_2_HPO_4_ 1. 2 g. L^−1^; KH_2_PO_4_ 1 g. L^−1^; hemin chloride (0. 1% in NaOH 0. 05 M) 10 ml. L^−1^; BSA (7. 5%) 24 ml. L^−1^; L-glutamine 4 g. L^−1^) at 28°C for one week.

### Artificially inoculated plant samples

For artificial inoculations, 10 ml of a calibrated CFBP 7970 or CFBP 8418 strain suspension was spiked in 2 g of detached *V. vinifera* leaves. Sterile water was used for negative controls. The DNA extraction was performed using a CTAB-based extraction protocol (2) with slight modifications in order to concentrate bacterial DNA. After a 20-min centrifugation at 20,000 g of the sample, the pellet was resuspended in 1ml of CTAB buffer. At the end of the extraction protocol, the pellet was resuspended in 50 μl of sterile demineralized water. *Xf* presence in the infected samples was checked using Harper’s qPCR assay (8).

### Plant material and bacterial gDNA

Healthy plant material of Vitis *vinifera (*2 g of leaf petioles) was spiked with 10 ml of a calibrated CFBP 7970 or CFBP 8418 strains suspension. Sterile water was used as negative control. The DNA extraction was performed using a CTAB-based extraction protocol (2) with slight modifications in order to concentrate bacterial DNA. After a 20-min centrifugation at 20,000 g of the plant macerate sample was resuspended in 1ml of CTAB buffer. At the end of the extraction protocol, the pellet was resuspended in 50 μl of sterile demineralized water. *Xf* presence in the infected samples was checked using Harper’s qPCR assay (8).

Naturally infected samples were collected in Europe and the USA. Symptomatic samples of *Olea europaea*, *Polygala myrtifolia* and *Quercus ilex* were collected in October 2018 in Corsica (France) and in September 2019 in the French Riviera. *Xf* detection and DNA extraction of whole infected-plant tissue were performed as mentioned above. *Xf* subspecies were identified using the tetraplex qPCR (17).

Twig tissues, leaf petioles or green shoots of *Rhamnus alaternus*, *Spartium junceum* and *P. myrtifolia* growing in the *Xf* outbreak zone of Monte Argentario (Grosseto, Tuscany, Italy) were collected during 2019 and 2020 (33). *Xf* was detected using Harper’s qPCR assay (8). For samples with a Cq value lower than 30, *Xf* isolation was attempted on Buffered Charcoal Yeast Extract (BCYE) agar according to PM 7/24-4 (2). Bacterial isolates that became visible to the unaided eye within three days of incubation at 28°C were discarded; those that became visible thereafter were streaked twice for purity on BCYE agar and identified as *Xf* based on qPCR results (8). Reactions were carried out after boiling bacterial suspension for 10 min. The DNA of one isolate among those that tested positive by qPCR from each plant, was extracted using the CTAB based protocol and further characterized to subspecies and ST level following the Multi Locus Sequence Typing (MLST) approach (34). The GoTaq probe qPCR Master Mix (Promega, A6102) and GoTaq G2 (Promega, M784B) polymerase were used for qPCR and conventional PCR experiments, respectively. The bacterial DNA of three isolates from *R. alaternus (*IT1-Ra to IT3-Ra), one from *S. junceum* (IT1-Sj), one from *P. myrtifolia* (IT1-Pm) and one *from Prunus dulcis* (IT1-Pd), were sequenced in this study. The assembled genomes of the six isolates were deposited on NCBI under the Bioproject PRJNA728043.

*Vitis vinifera* DNA samples were received from the Virginia Tech Plant Disease Clinic (VA, USA). All samples were collected from Virginia vineyards in 2019. US2-Vv sample was collected from the vineyard in Greene County and US3-Vv sample was from a vineyard in Isle of Wight County. The DNA extraction and *Xf* detection protocols are based on work instructions from the Virginia Tech Plant Disease Clinic (VTPDC). Approximately 50-100 mg of grape leaf or petiole tissue was excised from each sample using a razor blade. The excised tissue was transferred to lysing Matrix A tubes (MP, 6910-500) and ground using a FastPrep 24 (MP, 116004500). DNA was extracted using ISOLATE II Plant DNA extraction kit (Bioline, 52070) following manufacturer’s recommendations and CTAB lysis buffer. For *Xf* detection, qPCR Harper’s was performed on the StepOnePlusTM system (Life Technologies, 4376600) with Sensi-FAST Probe Hi-ROX qPCR kit (Bioline, 82005) (8).

### Pipeline controls

To test the pipeline, seven samples were used as negative controls. The controls were two DNA samples from healthy barley and wild grass leaves that were grown in a greenhouse, two DNA samples from barley leaves were infiltrated with a bacterial suspension (10^8^ CFU/ml) of *Xanthomonas translucens* pv. *translucens* UPB886. Three healthy samples were collected in France in 2020 and in the USA in 2019. Petioles and midribs were collected from healthy *Olea europaea* plants in a non-*Xf* infected area (Angers, France) and from *Polygala myrtifolia* plants that were purchased form a local nursery. *V. vinifera* leaves were collected from the vineyard in Greene County (Va, USA).The DNA from these samples was extracted using CTAB method as mentioned above. The absence of *Xf* in the healthy plants was confirmed using Harper’s qPCR assay (8).

For *in-silico* pipeline controls, two types of positive controls were used. To validate the detection of different concentrations of *Xf,* barley fasta sequence files were *in-silico* mixed with reads of the *Xff* CFBP 7970 sequenced in this study. The final proportion of *Xff* reads in the sample ranged from 0.2 to 2.4% of total reads. To validate the detection limits for single and mixed infections, the barley fasta sequence files were *in silico* mixed with different proportion of *Xff* CFBP 7970 and *Xfm* CFBP 8418 reads, to get from % CFBP7970: 99% CFBP8418 to 99% CFBP7970: 1% CFBP8418.

### iSeq 100 sequencing

iSeq 100 sequencing libraries were prepared according to the Illumina reference guide for Nextera DNA Flex Library Prep and Nextera DNA CD Indexes. In brief, 200 to 500 ng of DNA were quantified by spectrophotometry and used for library preparation. Then, the libraries were diluted to have the same starting concentration prior to sample pooling. Eight to 12 libraries were mixed together, and 1 nM of pooled library was used per run. The sequencing settings were paired-ended (PE) read type, 151 read cycles and eight index cycles. In the iSeq 100 System, the illumina GenerateFASTQ Analysis Module for base calling and demultiplexing was selected. After sequencing for 17 h, fastq paired end read files were extracted from the machine for subsequent analysis.

### Pipeline for *Xylella* sp. detection, classification and quantification via metagenomic analysis

Fastq files and the program Kraken2 were used to recover *Xylella* reads (22). The Kraken2 command options were: --paired, --minimum-hit-groups 5, --report and –db. The database was created with 92 NCBI genomes: 79 from *Xf*, two from *Xylella taiwanensis*, ten from *Xanthomonas* sp. and one from *Escherichia coli* (Table S1). The 81 *Xylella* genomes were used to recover *Xylella* reads. The last 11 genomes were added to remove reads common to Proteobacteria that might give a false positive match. The tool SendSketch with nt server was used to make sure the 81 NCBI *Xf* genomes were not contaminated with plant reads (BBMap – Bushnell B. – sourceforge.net/projects/bbmap/; October 30, 2019). Forty-nine NCBI sequences from plant 18S and chloroplast were added to the Kraken2 customized database to avoid extracting reads annotated as plant reads (Table S1).

The reads classified as *Xylella* were extracted with the script extract_kraken_reads from the KrakenTools suite (GitHub jenniferlu717/KrakenTools). The extracted *Xf* reads were used for downstream analysis. First *Xf* reads were *de-novo* assembled with the software SPAdes (23) using defaults settings and the option --only-assembler. Second, the *Xf* contigs were the query sequences in the Basic Local Alignment Search Tool website (NCBI) to confirm if they were *Xf* reads, or misclassified plant reads. The Blastn parameters were Nucleotide collection (nr/nt) as database and megablast program selection.

The *Xf*-positive contigs were used in four different analyses: 1) to identify subspecies, 2) to reconstruct phylogeny, 3) to identify the genetically closest strains already sequenced and 4) to identify alleles from specific genes and the MLST profile.

To identify subspecies, the tool SendSketch was run with the parameters, mode=sequence, records=2, and format=3, minani=100, minhit=1, address=ref, level=0 (BBMap – Bushnell B. – sourceforge.net/projects/bbmap/). Only contigs with 100% average nucleotide identity (ANI) were used to identify *Xf* subspecies. The *Xf* contigs with no hits were considered as core sequences. For visualization, results were plotted using stacked bars.

To reconstruct phylogeny, ANI was calculated using the software Pyani (v. 0. 2. 10) and LINbase (24, 25). For Pyani, the option -ANIm was set and for LINbase the “Identify using a gene sequence” was set as identification method. The R package ComplexHeatmap was used to visualize the ANI cluster analysis with the parameters clustering_distance_rows = robust_dist and clustering_method_rows = “average”. Robust_dist was a function suggested by the ComplexHeatmap tutorial.

To identify the genetically closest already sequenced *Xf* strains, the tool BBSplit was run with *Xf*-contigs (BBMap – Bushnell B. – sourceforge.net/projects/bbmap/). The tool used 81 *Xf* genomes from NCBI and the options minratio=1 ambig=best. For comparisons, each sample was normalized to their total *Xf* contigs and plotted using the R package ComplexHeatmap. For the isolated bacterial genomes, the most abundant strains were used as reference to identify genomics variants using Harvest suite tools (35).

To determine specific gene alleles, percentage of identity was calculated using *Xf* contigs as query and the Blastn algorithm (Nucleotide-Nucleotide BLAST 2. 8. 1+). The database contained complete nucleotide sequences for all alleles for the seven genes used for ST identification (*cysG, gltT, leuA, malF, nuoL, holC and petC*) (34). All 147 alleles were downloaded from the website PubMLST (36) (Last updated: 2019-03-06). To determine the presence of reported *Xf* virulence-related or common to several plant pathogenic bacterial genes (37, 38), percentage of similarity was calculated using *Xf* contigs as query using Blastx algorithm (Nucleotide-Nucleotide BLAST 2. 8. 1+. The databases contained complete amino acid sequences for gumBCDE, pilBMQTVW, rpfCEFG, tolC, 6-phosphogluconolactonase (pgl), and hemagglutinin from the *Xfm* M12 and *Xff* M23 NCBI genomes.

### MiSeq sequencing

To validate the iSeq 100 results, the same iSeq 100 libraries were used for MiSeq deep sequencing. Nine samples were selected, at least one from each iSeq 100 run, making sure not to use the same i5 and i7 tags. These nine libraries were sent to the Animal Disease Diagnostic Laboratory (Ohio Department of Agriculture, Reynoldsburg, Ohio) for sequencing. Library preparation was performed using an Illumina DNA Flex kit, and 2×250 sequencing was performed on the MiSeq platform using V3 chemistry. The pipeline described above was used to analyze the MiSeq fastq files.

## ACKNOWLEDGEMENTS

The authors are grateful for funding support from the Ohio Department of Agriculture Specialty Crops Block Grant (AGR-SCG-19-03) and USDA NIFA FACT (2021-67021-34343) to JMJ. Sara Campigli was financed by a grant from the Phytosanitary Service of the Tuscany Region (Italy). We would like to thank The Ohio Supercomputer Center for providing High Performance Computing resources.

We would also like to thank Dr. Stephen Cohen, Dr. Jeff Chang, and Dr. Alexandra Weinsberg for their valuable input for the experiment development and editing.

